# One New Species of the Genus *Dryomys* (Rodentia, Glirgae) from Xinjiang China, *Dryomys Yarkandensis sp*. nov

**DOI:** 10.1101/2020.02.11.943381

**Authors:** L. Liao, R. An, S. Shi, Y. Xu, Y. Luo, W. Liao

## Abstract

During an expedition in June 2012 in Shache county of Tarim Basin in southern Xinjiang, China, a new species of the genus *Dryomys* (Gliridae) has been discovered and named *Dryomys yarkandensis sp.* nov. It has been found obviously different from *D. nitedulai* in northern Xinjiang, *D. laniger, D. niethammeri* and *D. nitedulai* in Europe, which are also belong to genus *Dryomys*. The new species *Dryomys yarkandensis sp.* nov is described below.

**Holotype:** No. N07, an adult female collected by Chen Zhenhai in June 2012, is deposited Center for disease prevention and control of Xinjiang (Xinjiang CDC). It was obtained from oasis orchard of desert in Tarim Basin (38°29’N, 77°32’E), 1211-1215 m.

**Genus character:** There is a dark chestnut round eye. The terminal of tail is club shape, covered with dense hairs, and cannot see the scale ring in external texture.

**Description of the species:** The eyes is large. The beard is long and the longest could reach 30 mm. The tail is thicker and slightly longer than body length about 10%. The terminal of the tail is fluffy. All the surface is covered with dense hairs.

**External figure:** The color of the new species on the back is lighter than that of *D. laniger, D. niethammeri* and *D. nitedulai* in *Dryomys*. The length of the tail is about 110% of the body length. The length of the ears is 12.8 mm, which is 15% shorter than the other three species of *Dryomys*.

**Skull and tooth:** The ratio between the length of audirory bullae and the breadth of auditory bullae is 1.66 (8.35/5.02), which is larger than the other three species of *D. nitedula*.

The habitats of the new species is harsh, drought and hot in summer but dry and cold in winter. The habitats of *D. nitedula* in mountain valley in northern Xinjiang is temperate, humidity and low temperature, and there are berries or orchard.

## Introduction

*Dryomys nitedula* (Pallas, 1779) is the mode species established the genus *Dryomys* of family Gliridae (Thomas, 1906). It distributes extensively in central and southeastern Europe (Kryštufek and Vohralik, 1994), northern and eastern Russia, Turkey, the Middle East, Afghanistan, Pakistan, central Asia (Ognev, 1940; Allen, 1940; Sisson and Grossman, 1950; Bosessneck and Driesch, 1976; Sokolov, 1977; Sludskii, 1977; Kryštufek, 1999; Airapetyants and Fokin, 2005), as well as eastward to Dzungarian Alatau, Sawuer mountain, altai mountains in China and Khovd river upstream in western Mongolia (Shiirevdamba et al., 1997). *D. nitedula* lives in a wide range of habitats, including broad-leaved, mixed and coniferous woodlands, as well as rocky areas, dwarf mountain woodland, evergreen shrubland and wood-steppe (Kryštufek, 1999).

More than half a century, it has been considered as a monotype genera. According to previous work (Ellerman and Morrison-Scoot, 1951), 20 names species or subspecies of the genus had been published by 1946. But many taxonomists thought they are synonyms of *D. nitedula* (ОГнев, 1947; Ellerman and Morrison-Scoot, 1951). In 1968, Felton and Storch considered that the specimen of forest dormice, which was collected from the Turkish Anatolia Rocky Mountains, is different from *D. nitedula* in terms of size and morphology characteristics, so they published a new species, *Dryomys laniger* (Felton and Storch, 1968). Later, *Dryomys niethammeri*, a new species of *Dryomys* collected from Pakistan’s baluchistan province, was described and compared the morphology with *D. nitedula* and *D. laniger* (Holden, 1996). The research conclusions were recognized by another work (Rossolimo et al., 2001). At present, there are three species of the genus *Dryomys, D nitedula, D. laniger* and *D. niethammeri*.

Up to now, there is only one species of the genus *Dryomys* in China, *D. nitedula*, which distributes in Tianshan mountain, Altai mountain, Dzungarian Alatau and Yili basin (Wang and Yang, 1983; Ma et al., 1987; Zhang, 1997). In 2012, we obtained some specimens of dormice from oasis orchard of desert in Tarim Basin of southern Xinjiang, China. The specimens are similar to *D. nitedula*, but has morphological differences from *D. nitedula*. In order to find out its classification status, we collected living specimens to breed in lab from Shache county in Tarim basin of southern Xinjiang and form Jinghe county of northern Xinjiang in 2013. We compared the specimens from the two areas from adult to larval according to morphological feature.

## Part 1 Study of morphology on the dormice from Tarim basin in China

### 1. Materials

Specimens of the dormice from Tarim basin: n=15.

Specimens of *D. nitedulai* from northern Xinjiang in China: n=11, 6 specimens from mountains of northern Xinjiang deposited in Institute of Zoology, Chinese Academy of Sciences, including one from Altai, one from Fukang, one form Manas, one from Nileke counties, two from Talbahatai mountain of Tacheng prefecture and five from Jinhe county deposited in Xinjiang CDC Herbarium.

*Dryomys nitedulai*: the data of 85 specimen are quoted from previous work (Holden, 1996).

*Dryomys laniger*: the data of 17 specimen are quoted from previous work (Felten and Storch, 1968).

*Dryomys niethammeri*: the data of 3 specimen are quoted from previous work (Holden, 1996).

### 2. Methods

#### Age criteria

Specimens were assigned to larvae, juveniles and adult age primarily according to the body length, degree of tooth wear and the time of actual breeding.

#### Sexual dimorphism

Krystufek (Krystufek, 1985) found no statistically significant differences of morphology between males and females.

#### Color comparison

We compared the color of specimens from Tarim basin and northern Xinjiang with different ages, areas and collecting seasons.

#### Morphologic Measurements

The measurement methods were described in previous work (Holden, 1996; Yang et al., 2005; Xia et al., 2006). Cranial and dental measurements were taken with dial calipers graduated to tenths of millimeters. Skin color and body weights were recorded by collectors on the original skin tags. Measurements are abbreviated in the text as follows:

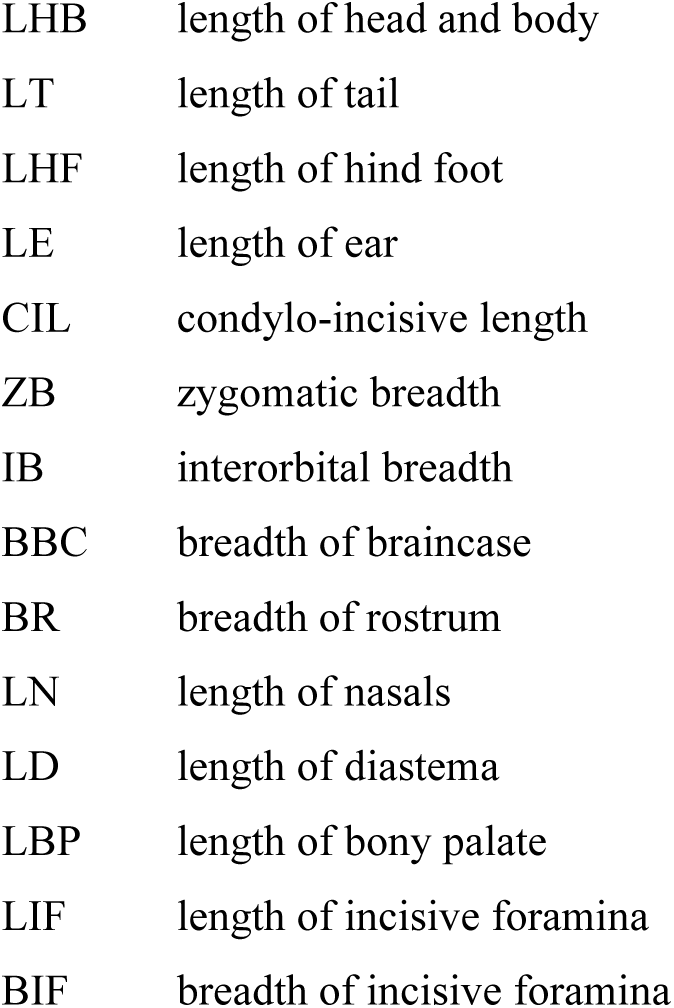
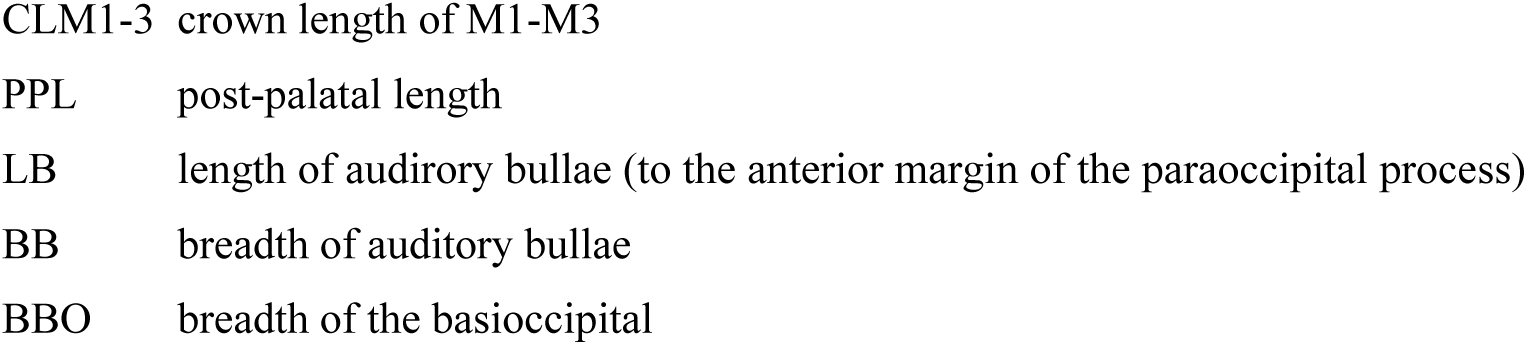

#### Statistical analysis

The measure data were analyzed by Origin7.5 (Data Analysis & Scientific Graphing Origin). Differences between the two groups of the specimens were tested by t-tests. Significant differences were considered according to the average value and standard deviation, in which *α* ≤ 0.05.

#### Ratio diagram

We used exactly the same method as Holden to make ratio diagram (Holden, 1996).

#### Principal components analysis

A principal components analysis was performed by Origin7.5 on variance-covariance matrix computed from seven log-transformed dimensions: length of body, length of tail, length of ear, length of incisive foramina, breadth of incisive foramina, length of audirory bullae and basioccipital breadth. These particular dimensions are chosen because they are informative and obtainable from most specimens. Other measurements available from the same set of specimens have relatively low scores and do not aid in separation of the samples in multivarate space when included (Holden, 1996).

#### Etymology

The species is named according to place where specimens were collected, i.e. Tarim (the largest basin in China).

## Part 2 *Dryomys yarkandensis sp*. nov from Tarim (a new species)

### 1. Information of the new species

#### Holotype

No. N07, an adult female collected by Zhenhai Chen in June 2012, is deposited Center for disease prevention and control of Xinjiang (Xinjiang CDC). It was obtained from oasis orchard of desert in Tarim Basin (38°29’N, 77°32’E), 1211-1215 m. The measurements are listed in table 1.

**Table 1.** The measurements of *Dryomys yarkandensis sp.* nov.

| No. | BW<br>(g) | LHB | LT | LE | LHT | CIL | LBP | LN | IB | ZB | BBO | LB | BB | BBC | LD | LIF |
| --- | --- | --- | --- | --- | --- | --- | --- | --- | --- | --- | --- | --- | --- | --- | --- | --- |
| Holotype N7 | 21.3 | 83 | 96 | 13 | 21 | 23.9 | 4.87 | 9.12 | 5.16 | 14.2 | 1.85 | 7.9 | 4.8 | 12.5 | 6.46 | 2.62 |
| Allotype N32 | 18.6 | 85 |  | 12 | 23 | 25.2 | 5.25 | 9.78 | 4.31 | 15.3 | 2.01 | 8.87 | 5.17 | 13.1 | 6.56 | 3.32 |
| Paratype N9 | 21.0 | 92 | 101 | 12 | 22 | 25.2 | 5.17 | 9.9 | 4.53 | 15.0 | 2.04 | 8.39 | 4.98 | 13.3 | 6.53 | 3.21 |
| Paratype N27 | 21.7 | 84 |  | 13 | 21 | 25.8 | 4.93 | 9.08 | 4.6 | 15.8 | 2.01 | 9.35 | 5.31 | 13.3 | 6.86 | 2.96 |
| Paratype N14 | 24.2 | 83 | 98 | 12 | 22 | 25.1 | 5.09 | 9.56 | 4.2 | 15.5 | 1.81 | 8.45 | 5.18 | 12.9 | 6.6 | 2.68 |
| Paratype N23 | 23.7 | 84 | 85 | 12 | 22 | 26.2 | 5.61 | 9.35 | 4.43 | 15.5 | 1.89 | 8.69 | 4.97 | 13.2 | 6.72 | 2.76 |
| Paratype N15 | 23.6 | 83 | 100 | 13 | 21 | 24.6 | 4.43 | 8.44 | 4.3 | 14.6 | 1.9 | 8.51 | 4.6 | 12.4 | 6.25 | 2.92 |
| Paratype N35 | 22.7 | 86 |  | 14 | 22 | 24.4 | 4.98 | 9.34 | 4.33 | 14.4 | 1.77 | 7.52 | 4.73 | 12.5 | 6.08 | 2.41 |
| Paratype N34 | 22.0 | 81 | 101 | 14 | 22 | 25.8 | 5.39 | 9.52 | 4.69 | 16.1 | 1.95 | 8.65 | 5.84 | 13.3 | 6.52 | 2.88 |
| Paratype N44 | 21.5 | 82 | 95 | 15 | 22 | 24.2 | 4.8 | 9.51 | 4.58 | 15.0 | 1.91 | 8.69 | 5.11 | 13.6 | 6.42 | 2.67 |
| Paratype N13 | 21.3 | 85 | 99 | 12 | 20 | 24.2 | 5.03 | 8.98 | 4.2 | 14.7 | 1.75 | 8.29 | 5.03 | 12.8 | 6.3 | 2.49 |
| Paratype N40 | 21.1 | 96 |  | 14 | 21 | 24.8 | 4.75 | 9.6 | 4.55 | 14.6 | 1.81 | 8.14 | 4.8 | 12.7 | 6.6 | 2.76 |
| Paratype N43 | 21.1 | 92 |  | 11 | 20 | 25.3 | 5.3 | 9.75 | 4.5 | 15.3 | 1.91 | 8.39 | 5.15 | 12.9 | 6.44 | 2.53 |
| Paratype N20 | 21.1 | 83 |  | 13 | 22 | 25.1 | 4.44 | 9.3 | 4.17 | 14.4 | 1.81 | 8.17 | 4.71 | 12.7 | 6.39 | 2.51 |
| Paratype N33 | 20.6 | 85 |  | 11 | 21 | 24.7 | 5.73 | 9.44 | 4.36 | 15.4 | 1.76 | 7.7 | 5.12 | 13.1 | 6.45 | 2.52 |
| X | 21.7 | 85.6 | 96.9 | 12.7 | 21.5 | 25.0 | 5.1 | 9.35 | 4.46 | 15.1 | 1.88 | 8.38 | 5.03 | 13.0 | 6.48 | 2.75 |
| SD | 1.41 | 4.29 | 5.28 | 1.16 | 0.83 | 0.66 | 0.38 | 0.36 | 0.25 | 0.56 | 0.10 | 0.46 | 0.30 | 0.35 | 0.19 | 0.27 |

#### Allotype

No. N32, an adult male deposited in Xinjiang CDC, was obtained from Tarim Basin in June 2012. The rest information is the same as holotype.

#### Paratype

No. N9, an adult female, is deposited in Institute of Zoology, Chinese Academy of Sciences. No. N27, an adult female, is deposited in Xinjiang CDC. The rest information is the same holitype.

The other 11 adult female specimens deposited in Xinjiang CDC, were obtained from Tarim Basin at the same time. The rest information was as same as holotype. All the measurement data are listed in table 1.

#### Contrast samples

Specimens of *D. nitedula* from northern Xinjiang, n=11.

Pups picture: 16-day-old and 26-day-old pups of both *D. nitedula* collected in Jinghe county and the new species collected in Shache county were all bred in laboratory (figure 1c,d,e,f). All the measurement data are listed in table 1,2.

**Table 2.** Compare of measurements between *D. yarkandensis sp* and *D. nitedula* from northern Xinjiang.

| Measurement | Average ± SD (n) mm | | | T-test | Difference<sup>#</sup> |
| --- | --- | --- | --- | --- | --- |
|  | Holotype N7 | *D. y* (14) | *D. n* (11) |  |  |
| Body weight (g) | 21.3 | 21.91 ± 1.166 | 26.4 ± 4.0 | $P = 0.0014^*$ | -13.92 |
| Head & body length | 83.6 | 85.64 ± 4.448 | 92.73 ± 6.51 | $P = 0.0063^*$ | -7.51 |
| Tail length | 96 | 96.88 ± 3.276 | 93.9 ± 8.61 | $P = 0.2319$ | 4.54 |
| Ear length | 13 | 12.79 ± 1.188 | 16.45 ± 2.69 | $P = 0.00014^{**}$ | -19.2481 |
| Hind foot length | 21 | 21.36 ± 0.745 | 20.27 ± 2.28 | $p = 0.2522$ | 2.68 |
| CIL | 23.9 | 24.94 ± 0.682 | 25.74 ± 1.14 | $P = 0.1352$ | -5.83 |
| Condylobasal length | 22.6 | 24.01 ± 0.775 | 24.67 ± 1.69 | $P = 0.2356$ | 0.99 |
| LBP | 4.9 | 5.04 ± 0.385 | 4.79 ± 0.182 | $P = 0.0839^{**}$ | -5.83 |
| LN | 9.1 | 9.35 ± 0.364 | 8.26 ± 0.65 | $P < 0.0002^{**}$ | 13.91 |
| IB | 5.2 | 4.47 ± 0.258 | 4.41 ± 0.157 | $P = 0.5347$ | 1.32 |
| ZB | 14.2 | 15.03 ± 0.590 | 16.04 ± 0.573 | $P = 0.0062^*$ | -6.25 |
| BBO | 1.9 | 1.87 ± 0.091 | 2.28 ± 0.334 | $P = 0.0006^{**}$ | -16.46 |
| LB | 7.9 | 8.35 ± 0.460 | 7.39 ± 0.541 | $P < 0.0001^{***}$ | 13.50 |
| BB | 4.8 | 5.02 ± 0.313 | 5.41 ± 0.295 | $P = 0.0191^*$ | -6.32 |
| BBC | 12.5 | 12.95 ± 0.378 | 13.34 ± 0.372 | $P = 0.0290^*$ | -2.76 |
| LD | 6.46 | 6.47 ± 0.195 | 6.41 ± 0.482 | $P = 0.4591$ | 1.68 |
| LIF | 2.62 | 2.71 ± 0.223 | 2.94 ± 0.371 | $P = 0.2024$ | -4.71 |
| BIF | 2.21 | 2.51 ± 0.205 | 1.88 ± 0.169 | $P < 0.0001^{***}$ | 34.99 |
| Interparietal length | 2.8 | 2.74 ± 0.37 | 3.37 ± 0.528 | $P = 0.475$ | -16.3 |
| Interparietal width | 8.55 | 8.58 ± 0.55 | 9.61 ± 0.717 | $P = 0.0023^*$ | -10.16 |
| Anterior palatal breadth | 3.6 | 3.9 ± 0.21 | 4.24 ± 0.402 | $P = 0.0023^*$ | -7.5 |
| Posterior palatal breadth | 3.6 | 3.81 ± 0.11 | 4.21 ± 0.35 | $P = 0.0277^*$ | -8.64 |

\*: $p < 0.05$ , \*\*: $p < 0.01$ , \*\*\*: $p < 0.001$ , #: $(D. y - D. n) \times 100 / D. n$ , SD: Standard deviation,
*D. y*: *D. yarkandensis* sp., *D. n*: *D. nitedula* in northern Xinjiang.

**Figure 1.**
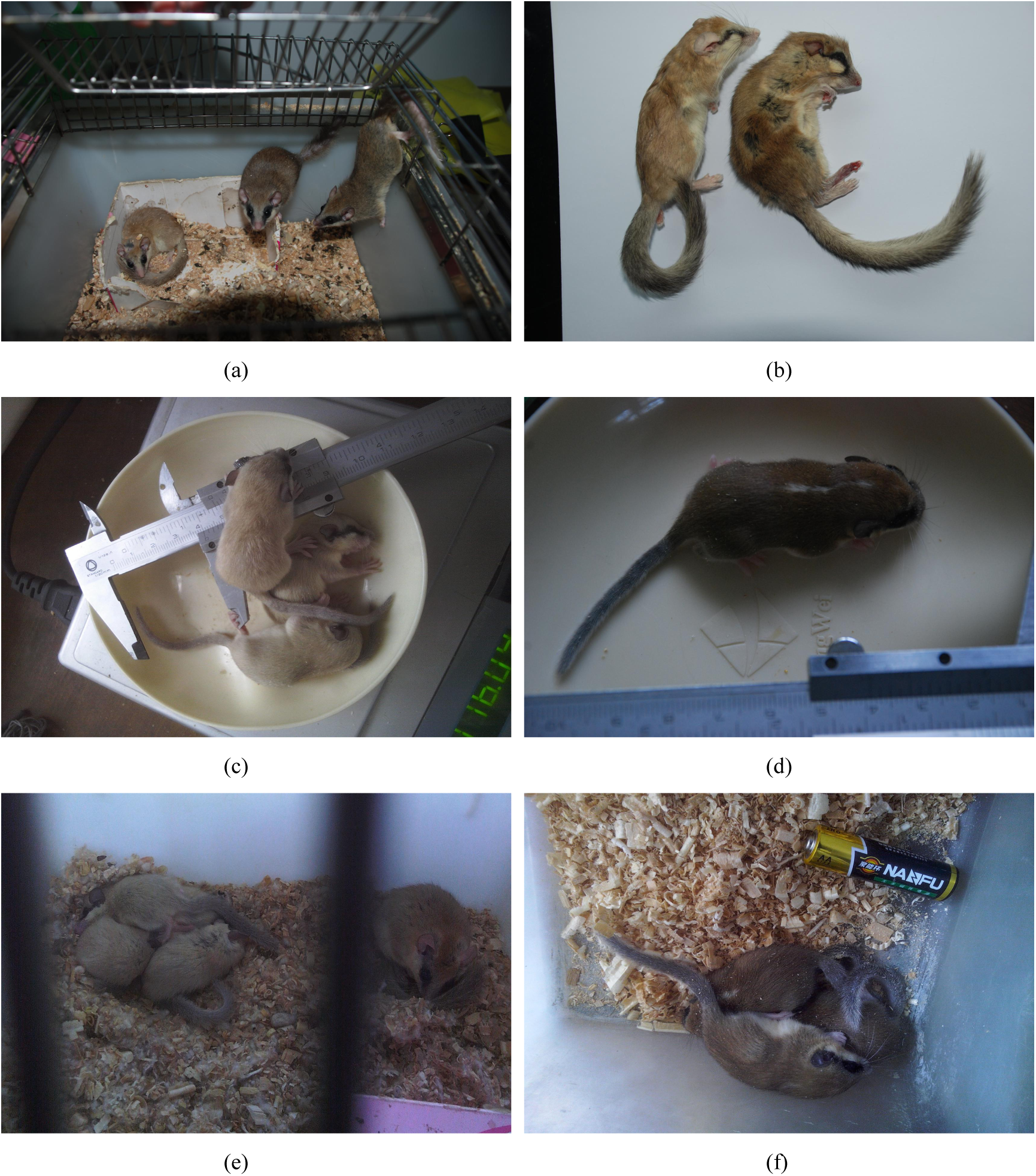
The coat color of larvae and adults of the new species and *D. nitedula* in northern Xinjiang. (a) The coat color of adults of the new species (the left one) and *D. nitedula* in northern Xinjiang (the right two). (b) The black stripe from ears across eyes in the new species (the left one) and *D. nitedula* in northern Xinjiang (the right one). (c) The 16-day larval of the new species. (d) The 16-day larval of *D. nitedula* in northern Xinjiang. (e) The 26-day larval of the new species. (f) The 26-day larval of *D. nitedula* in northern Xinjiang.

### 2. Description of holotype of the new species

#### Appearance

Its appearance is squirrel-like. Coat on the body and tail is heavy and soft, on the caudal is longer. There is a light black stripe from ear root through the eye to nose on both sides of the head each (figure 1a,b). The range of its body mass is between 16 grams and 30 grams. The average length of body is 85.6 (81-96) mm and length of the tail is 96.88 (85-101) mm, which is 110% of the body length. Ears short and rounded, its average length is 12.8 (11-15) mm. The length of hind foot is 21.4 (20-22) mm. There are four toes on the fore foot and five toes on the hind foot, on which there are six pads. The bilateral symmetry whiskers form a bushy tuft about 25-30 mm long.

#### Physical description

The back color of adult is hazel in Summer, and turn light brown after Autumn gradually, which close to tail color. Hair tip of belly is cream yellow, and root hair of belly is white. There is the boundary line between side of the body and belly, and the color of tail is dark than the back, light brown, and abdominal hair of tail slightly shallow.

There is light chestnut from the head to back of pup between 16-day-old and 26-day-old after birth, and a white on belly. At autumn the color of back and tail are turn light brown, hair tip of belly become cream yellow. Also the color of tail is dark than back and the boundary line between side of the body and belly become clear (figure 1a,c,e).

#### Skull characteristic

There are two pairs of incisors, smaller incisive foramina, developed heekbones, bulky zygomatic arch, bigger auditory bullae, which is separated into several small space by bone membrane on the kull. The dental formula: 1:0:1:3/1:0:1:3. The chewing surfaces of each molar has a few columns of transvrsal enamel teeth ridge. The length of incisive foramina is very close to length of interparietal bone (or incae), which is 11% of CIL. The ratio of LB to BB is 1.66 (8.35/5.02), and the length of audirory bullae is 2.28 (8.35/3.66) times of upper toothrow length. Junction between interparietal bone and two parietal bone is 130 degrees of a contact angle (figure 2b). The nasal bone embed into frontal bone with a convex shape (figure 2d).

**Figure 2.**
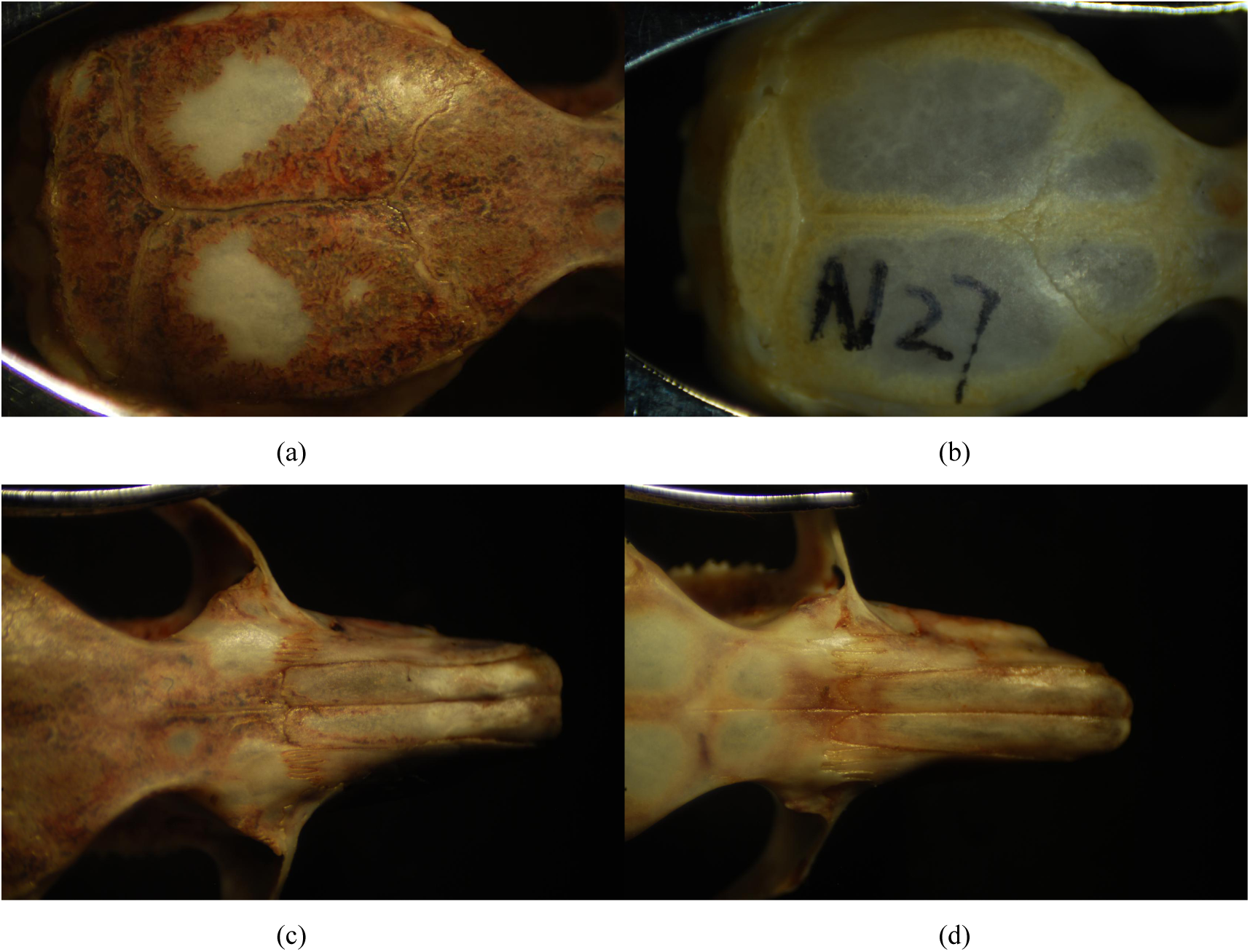
Comparison of the nasals and interpariet bone between the new species and *D. nitedula* in northern Xinjiang. (a) Interpariet bone of *D. nitedula* in northern Xinjiang. (b) Interpariet bone of the new species. (c) The nasals of *D. nitedula* in northern Xinjiang. (d) The nasals of the new species.

#### Skull Measurements

The measurement data are shown in Table 1. Basicranial Length: 24.9 (23.9-25.4) mm. Interorbital Breadth: 15 (14,4 −16.1) mm. Length of diastema: 9.35 (8.4-9.9) mm. Breadth of the basioccipital: 1.87 (1.8-4.04) mm. Length of audirory bullae: 8.35 (4.6-5.2) mm. Upper Toothrow Length: 3.66 (3.53-3.79) mm. Diameter of M3: 1.05 (0.98-1.1) mm. Breadth of incisive foramina: 2.51 (2.2-2.86) mm. Length of bony palate: 5.04 (4.43-5.61) mm. Breadth of braincase: 12.94 (12.4-13.6) mm. Height of braincase: 11.13 (10.85-12.1) mm.

### 3. Identification features of the new species

By comparing morphology (table 1 and figure 1,2) and statistical analysis (table 2,3), there are differences in geography distribution (figure 3,4) and habitat among the dormice from Tarim basin and the specimens from *D. nitedulai, D. laniger* and *D. niethammeri*. The dormouse from Tarim basin is considered as a new species of the genus *Dryomys* and named *Dryomys yarkandensis sp.*.

**Table 3.** The measurement values on the new species and the other three species of the genus *Dryomys*.

| Measurement items | Average±Standard deviation (n) mm |  |  |  |  |
| --- | --- | --- | --- | --- | --- |
|  | *D. niethammeri* | *D. nitedula* | *D. laniger* | *D. nitedula* in X.J | The new species |
| Body weight (g) | 33 (1) | 28.9 ± 7.1 (11) | 22.2 ± 3.73(17) | 26.4 ± 4.0 (8) | 21.91 ± 1.17 (14) |
| Head & body length | 101 ± 2.82 (3) | 97.2 ± 9.7 (55) | 89.9 ± 3.96(17) | 92.73 ± 6.51(11) | 85.64 ± 4.45 (14) |
| Tail length | 93 ± 0 (1) | 87.7 ± 11.88(44) | 66.9 ± 6.15(15) | 93.9 ± 8.61 (10) | 96.88 ± 3.28 (8) |
| Ear length | 19.9 ± 1.41 (2) | 14.9 ± 2.38 (52) | 14.4 ± 1.38(16) | 16.45 ± 2.69(10) | 12.79 ± 1.19 (14) |
| Hind foot length | 21 ± 0 (2) | 21 ± 1.51 (55) | 16.8 ± 1.1 (17) | 20.27 ± 2.28(11) | 21.36 ± 0.75 (14) |
| CIL | 25.2 ± 0.26 (3) | 23.9 ± 1.15 (56) | 23.1 ± 3.73(17) | 25.74 ± 1.14 (11) | 24.94 ± 0.68 (14) |
| LBP | 5.2 ± 0.07 (2) | 5.3 ± 0.31 (37) | 5.2 ± 0.35 (15) | 4.79 ± 0.182 (9) | 5.04 ± 0.385 (14) |
| LN | 9.3 ± 0.26 (3) | 9.1 ± 0.44 (36) | 8.4 ± 0.47 (13) | 8.26 ± 0.65 (11) | 9.35 ± 0.36 (14) |
| IB | $3.9 \pm 0.15$ (3) | $4.1 \pm 0.19$ (39) | $4.2 \pm 0.16$ (18) | $4.41 \pm 0.16$ (11) | $4.47 \pm 0.26$ (14) |
| ZB | $15.9 \pm 0$ (1) | $15.7 \pm 0.85$ (30) | $14.4 \pm 0.47$ (13) | $16.04 \pm 0.57$ (5) | $15.03 \pm 0.59$ (14) |
| BBO | $1.4 \pm 0.1$ (3) | $2 \pm 0.26$ (57) | $1.4 \pm 0.16$ (16) | $2.28 \pm 0.33$ (9) | $1.87 \pm 0.09$ (14) |
| LB | $9 \pm 0.15$ (3) | $7.3 \pm 0.45$ (60) | $8.1 \pm 0.21$ (18) | $7.39 \pm 0.54$ (11) | $8.35 \pm 0.46$ (14) |
| BB | $5.9 \pm 0.15$ (3) | $5 \pm 0.29$ (55) | $5.3 \pm 0.18$ (17) | $5.41 \pm 0.30$ (9) | $5.02 \pm 0.31$ (14) |
| LB/BB | 1.53 | 1.46 | 1.53 | 1.37 | 1.66 |
| BBC | $12.9 \pm 0.75$ (3) | $12.8 \pm 0.39$ (37) | $12.6 \pm 0.26$ (12) | $13.34 \pm 0.37$ (11) | $12.95 \pm 0.38$ (14) |
| LD | $6.3 \pm 0.2$ (3) | $6.3 \pm 0.34$ (39) | $5.8 \pm 0.25$ (18) | $6.41 \pm 0.48$ (11) | $6.47 \pm 0.20$ (14) |
| LIF | $3.9 \pm 0.06$ (3) | $3.6 \pm 0.29$ (39) | $3.4 \pm 0.2$ (17) | $2.94 \pm 0.37$ (9) | $2.71 \pm 0.22$ (14) |
| BIF | $2.2 \pm 0.06$ (3) | $2 \pm 0.19$ (39) | $1.9 \pm 0.11$ (18) | $1.88 \pm 0.17$ (7) | $2.51 \pm 0.21$ (14) |

**Figure 3.**
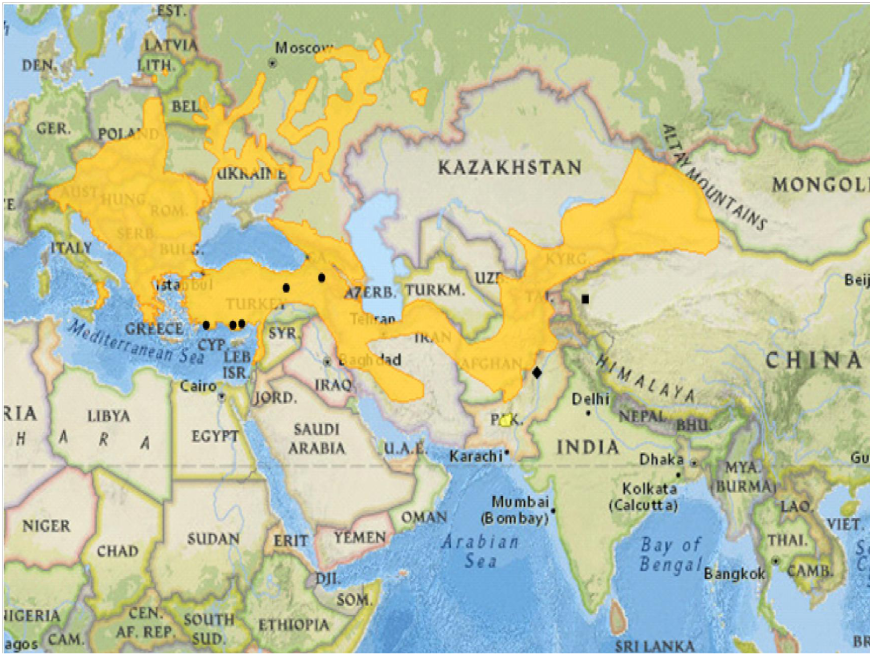
Distribution of different species of the genus *Dryomys* in Eurasia. The yellow color and the symbol ●, ◆, ■ represent *D. nitedula, D. laniger, D. niethammeri* and the new species *D. yarkandensis sp.* respectively.

**Figure 4.**
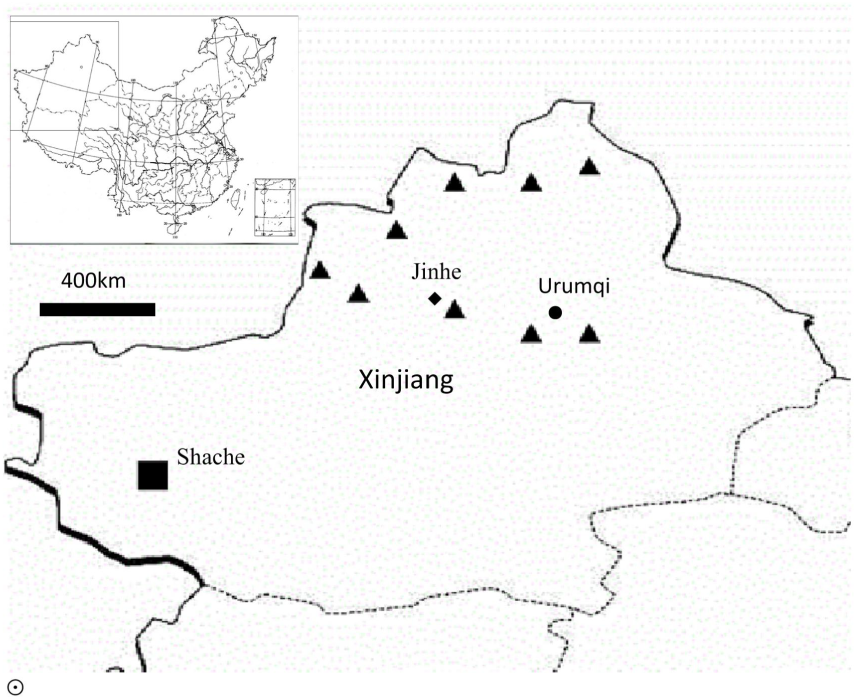
Distribution of the new species *D. yarkandensis sp.* (■) and *D. nitedula* (▲) of northern Xinjiang in China.

The skull shape of the specimens from Tarim Basin is similar to Muridae, but has a smaller incisive foramina and bigger auditory bullae, which is divided into several small space by bone membrane. There have well-developed cheekbones, thick zygomatic arch and two pairs of incisors on the skull. Dental formula: 1:0:1:3/1:0:1:3. The chewing surfaces of each molar have several columns of transvrsal enamel teeth ridge.

The color of pup is light chestnut from the head to the back (figure 1c,e), and a white on the belly. At autumn the color of back and tail are gradually turn light brown, hair tip of belly become cream yellow, also the color of tail is dark than back and the boundary line between side of the body and belly become clear. The length of incisive foramina is very close to length of Interparietal bone (or incae).

The ratio between length of audirory bullae and breadth of auditory bullae is 1.66 (8.35/5.02), and the length of audirory bullae is 2.28 (8.35/3.66) times of Upper Toothrow Length. Junction between Interparietal bone and two parietal bone is 130 degrees of a contact angle (figure 2b).The nasal bone embed into frontal bone with a convex shape (figure 2d).

## Part 3 Comparison with other three species of the genus *Dryomys*

### 1. Comparison with *D. nitedula* in northern Xinjiang

#### 1.1 Physical description

##### Specimens from Tarim basin

The color of adult on the back is lighter hazel. The color of pup or sub-adult is light chestnut hair from head to back, and a white on belly. There are a light black stripe from ear root through the eye to nose on both sides of the head each (figure 2). The length of the tail is significantly longer than body, more than 10% of the body length (table 1).

##### *D. nitedula* from northern Xinjiang

Coat on the body of the adult, pup and sub-adult are russet. The black stripe is darker from ear root through the eye to nose on both sides of the head. The length of tail is equal or less to the length of the body.

#### 1.2 Cmparison of skull measurements with *D. nitedula* from northern Xinjiang

There are ten indexes of skull measurements in the new species very close to *Dryomys* in northern Xinjiang, which was not statistically significant (table 2). There are very significant difference in 7 mean values of LN, BBO, LB, UTL, DM3, BIF, LBP and significant difference in the mean value of LN, ZB, BB, BBC, Interparietal width, Anterior palatal bread, and Posterior palatal bread, in which there are four index value (DM3, BBO, Interparietal width and UTL) of new species over 10% of *Dryomys* in northern Xinjiang, and three index value (LN, LB and BIF) less than 11% of *D. nitedula* in northern Xinjiang. The ratio of LB to BB in new species is 1.66 higher than the ratio (1.53) of *D. nitedula* in northern xinjiang. The ratio of LB to length of upper toothrow in the new species is 2.28, which is large than the ratio in *D. nitedula* of northern Xinjiang (1.83). The length of incisive foramina is close to the length of interparietal bone in the new species, but only 88% in *D. nitedula* of northern Xinjiang.

#### 1.3 Skull and teeth

Junction between interparietal bone and two parietal bone is 130 degrees of a contact angle in the new species (figure 2b), but less than 70 degree contact angle for *D.nitedula* of northern Xinjiang (figure 2a). The nasal bone embed into frontal bone with a convex shape for the new species (figure 2d), but with a flat mouthconvex shape for *D. nitedulai* in northern Xinjiang (figure 2c). There are open angle of two odontoid processes in third molar of maxillary (upper jaw) and four slots of the premolar in the new species (figure 5a), while amall open angle of two odontoid processes in third molar of maxillary (upper jaw) and three slots of the premolar in *D. nitedula* of northern Xinjiang (figure 5c). The transverse diameter of the third molar on mandibular was significantly smaller than that of the second molar in the new species (figure 5b), while the transverse diameter of the third molar on mandibular was slightly smaller than that of the second molar in *D. nitedula* of northern Xinjiang (figure 5d). The differences of the maxillary of skulls between the two species are shown in table 1-3, and the differences in appearance are shown in figure 6.

**Figure 5.**
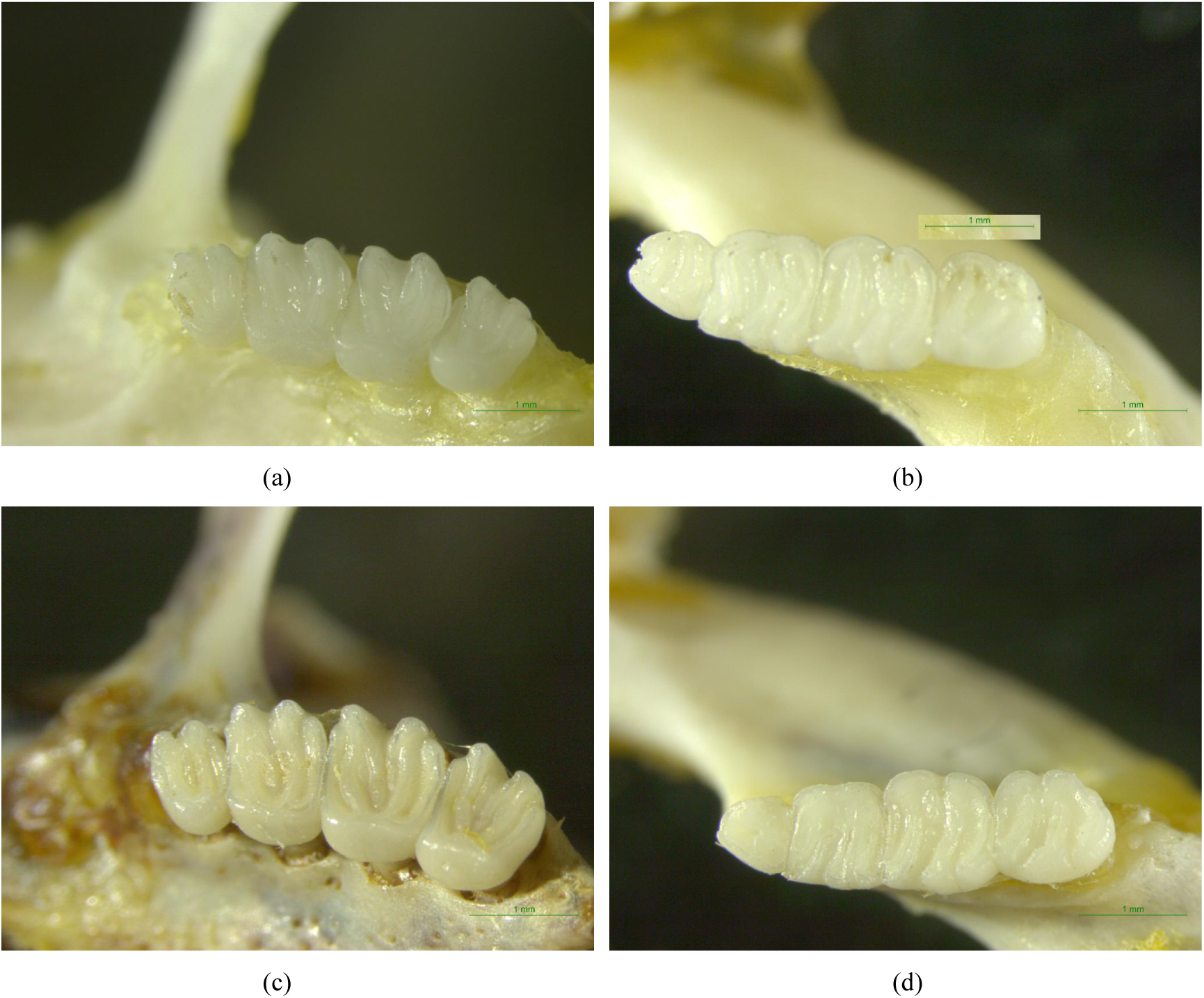
Comparisons of the crowns between the new species and *D. nitedula* in northern Xinjiang. (a) The dentition on left side of maxillary in the new species. (b) The dentition on left side of mandible in the new species. (c) The dentition on left side of maxillary in *D. nitedula* in northern Xinjiang. (d) The dentition on left side of mandible in *D. nitedula* in northern Xinjiang.

**Figure 6.**
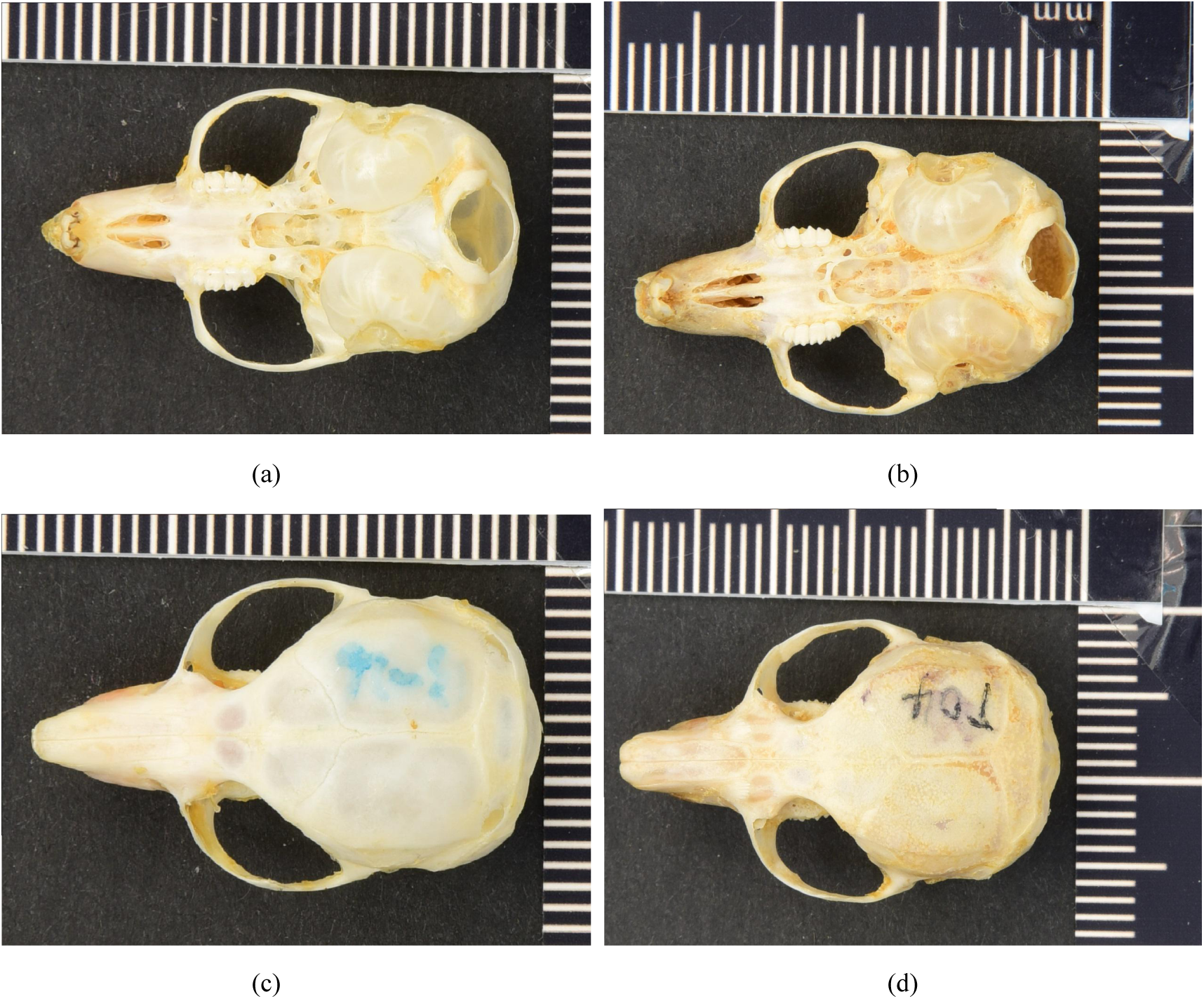
Comparison of the skulls between the new species and *D. nitedula* in northern Xinjiang. T04 is *D. nitedula* in northern Xinjiang and N07 is the new species *D. yarkandensis sp.*. (a) Ventral view of maxillaryfor *D. yarkandensis sp.* (b) Ventral view of maxillary for *D. nitedula.* (c) Dorsal view of maxillary for *D. yarkandensis sp.* (d) Dorsal view of maxillary for *D. nitedula.*

#### 1.4 Habitat comparison

The new species is distributed in the desert oasis orchard, 1200 m above sea level in Tarim Basin. The climate there is harsh, drought and hot in summer but dry and cold in winter (figure 7a). *D. nitedula* in northern Xinjiang is mainly distributed in mountain valley between 700 m to 1200 m above the sea level. The climate there is temperate, humidity and low temperature, and there are berries or orchard (figure 8b). It is obvious difference for the climates and habitats between the two distributions.

**Figure 7.**
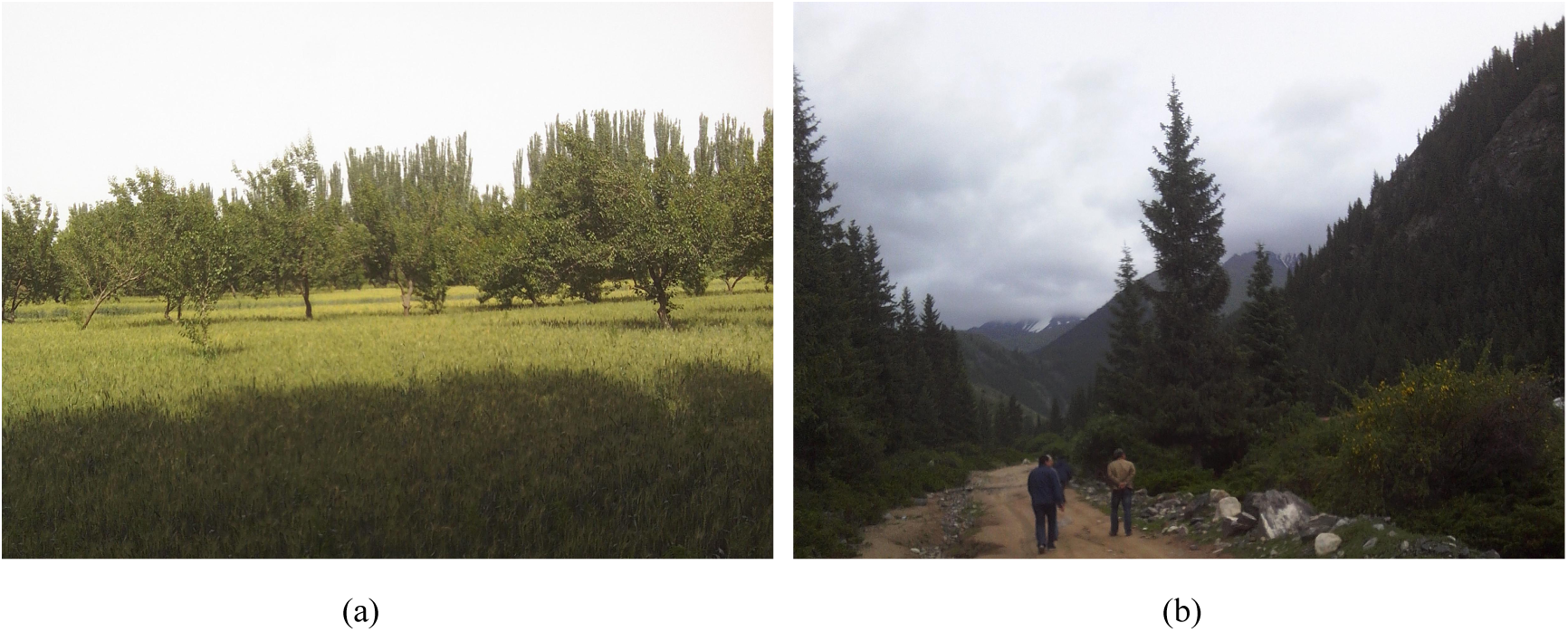
Habitat of (a) the new species and (b) *D. nitedula* in northern Xinjiang.

**Figure 8.**
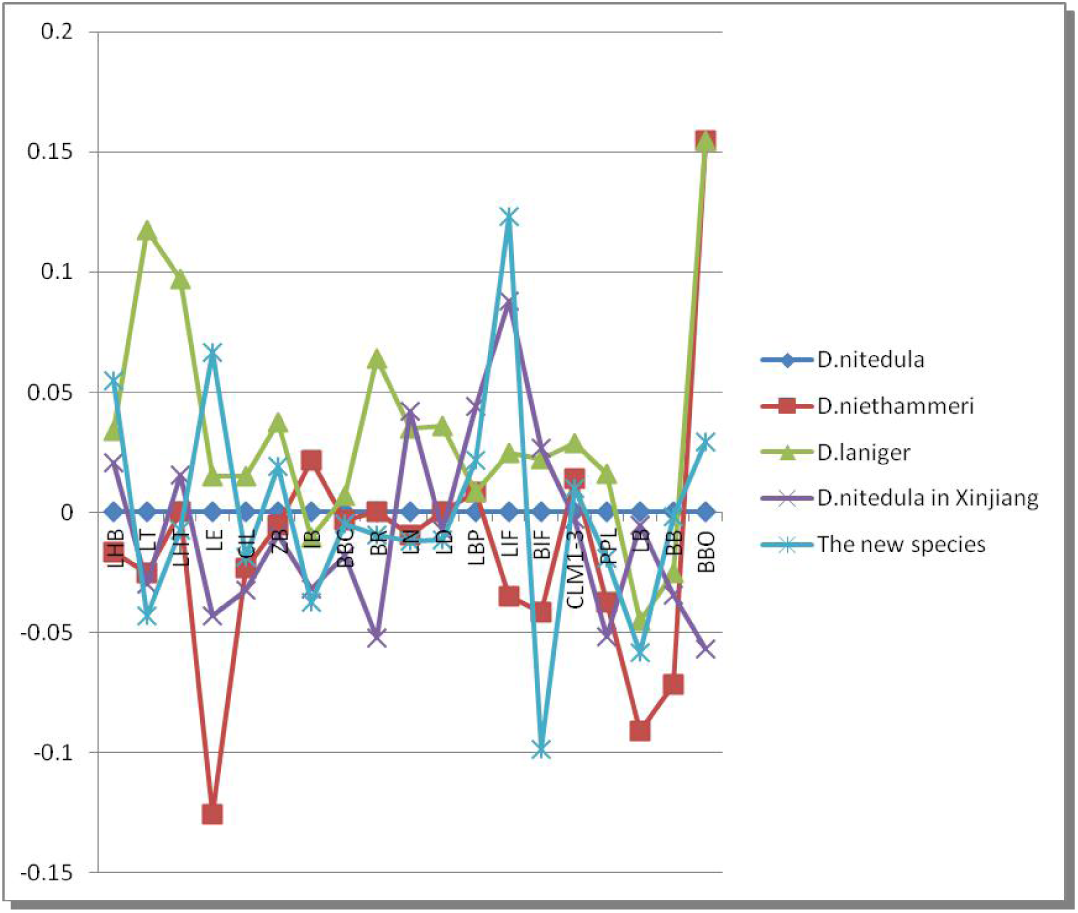
Comparison of the measured values between the new species and other species of *Dryomye*.

The color over the back of a new species is similar to yellow sand of the habitat backgrounds. The color of *D. nitedula* in northern Xinjiang is similar to brown of coniferous forest or dwarf shrub of the mountains in northern Xinjiang. This significant differences in color protection takes the dormouse to survive in them habitats.

### 2. Comparison with other three species in *Dryomye*

#### 2.1 Comparison of morphology

The adult of new species displays a lighter color of bodily hair on the back than that of other three species of *Dryomye*, and the black spot, which spread on both sides of head from the rear of nose, through eyes, then to the front of ears, is lighter-color than that of other three species of *Dryomye*. The LT of a new species obviously exceeds its LHB by 10%, however, the LTs of the other three species in Dryomye are less than or equal to body length (table 2,3).

#### 2.2 Comparison of skull measurements

As shown in figure 8 and table 3, the main different features between the new species and *D. laniger* are focused on LT, BIF, and BBO. Accordingly, the difference related to *D. niethammeri* and *D. nitedulai* are focused on LE, LIF, BBO and LHB, LT, LE, LIF, LB, BB, BBO, respectively. The ratio of LB to BB in the new species (1.66) is apparently higher than 1.40 in *D. laniger*, 1.53 in *D. niethammeri* and 1.46 in *D. nitedulai*. The ratio of LB to LD is 1.29 in the new species, while 1.4 in *D. laniger*, 1.43 in *D. niethammeri* and 1.16 in *D. nitedulai*.

### 3. Classified discussion

According to the observed value of morphology and the skull measurements, the new species of Tarim Basin differs significantly in morphological from *D. nitedulai, D. laniger* and *D. niethammeri*. Also considering the results of differences among habitats, we identified that the species of Tarim Basin is a new species, and its molecular genetics identification is going to further research. As a new species, it still remains empty for its distribution range, population situation and biological characteristics. It is urgent to develop further research on the new species from viewpoint of protection biology.

#### Distribution

The new species specimens had been obtained from this point in Tarim Basin only at present, other areas did not investigate in Xinjiang (figure 7b).

#### Biology data

Missing.

#### Check samples

11 specimen of *D. nitedula* from northern Xinjiang, 6 deposited in Institute of Zoology, Chinese Academy of Sciences, 21247 ♂ in Altai county, 21752 ♀ in Fukan county, 26659 ♀ in Manashi county, 26708 ♀ in Nileke county, 27482 ♀ and 27481 ♀ in Sawuer mountain; 5 specimens deposited in the Herbarium of Xinjiang CDC, T3 ♀, T4 ♀, T5 ♂, T31 ♀and T32 ♂ from Jinghe county of northern Xinjiang.

Reference sample data: from previous work (Holden, 1996).

*D. niethammeri*: n=3, Pakistan, 3 ♀.

*D. laniger*: n =21, Turkey, 5 ♂, 16 ♀.

*D. nitedulai*:n=85, Afghanistan n=10 (5♀, 5♂), Austria 2♀, Albania 1♀, Croatia n=2 (1 ♂, 1?), Georgia1?, Greece1♀, Hungary, n=2 (1♀, 1♂), Iran n=13 (6♀, 1♂, 6?), Italy1♂, Kazakhstan2♂, Lebanon 1?, Pakistan, n=9 (6♂, 3♀), Israel1♀, Poland2♂, Romanian=5 (3♀, 1♂, 1?), Russia2♀, Sweden1♂, Turkey n=23 (11♀, 10♂, 2?), Former-Yugoslavia, n=7 (3♀, 3♂, 1?).

## Acknowledgment

First of all, we would like to thank Professor Yong MA in Institute of Zoology, the Chinese Academy of Sciences, for kind guidance on this article. For permission to conduct research in Xinjiang, we would like to thank Zhenghai CHENG for his contributions to information and collecting samples from Tarim Basin. The data were measured and the samples were raised in our lab with the help of students, especially Xinhui LI and Xiaoyan LIAO in Xingjiang Agricultural University.

